# Studies of infused megakaryocytes into mice support a “catch-and-release” model of pulmonary-centric thrombopoiesis

**DOI:** 10.1101/2024.06.04.597316

**Authors:** Hyunjun Kim, Danuta Jarocha, Ian Johnson, Hyunsook Ahn, Nicholas Hlinka, Deborah L. French, Lubica Rauova, Kiwon Lee, Mortimer Poncz

**Author notes:** Correspondence: Kiwon Lee, Ph.D., Hankuk University of Foreign Studies Yongin, South Korea.

## Abstract

Many aspects of thrombopoiesis, the release of platelets from megakaryocytes (Mks), remain under debate, including where this process occurs. Murine lung *in situ*-microscopy studies suggested that a significant fraction of circulating platelets was released from lung-entrapped, marrow-derived Mks. We now confirm these that endogenous murine (m) Mks are immobilized in the lungs. Intravenously infused *in vitro*-differentiated, mMks and human (h) Mks are similarly entrapped, followed by shedding of their cytoplasm over ∼30 minutes with a peak number of released platelets occurring 1.5-4 hours later. However, while infused Mks from both species shed large intrapulmonary cytoplasmic fragments that underwent further processing into platelet-sized fragments, the two differed: many mMks escaped from and then recycled back to the lungs, while most hMks were enucleated upon first intrapulmonary passage. Infused immature hMks, inflammatory hMks, umbilical cord-blood-derived hMks and immortalized Mk progenitor cell (imMKCL)-derived hMks were also immobilized in the lung of recipient mice, and released their cytoplasm, but did so to different degrees. Pharmacologically-treated hMks support that membrane stiffness was key to pulmonary thrombopoiesis. Intraarterial infused hMks resulted in few Mks being immobilized in tissues other than the lungs and was accompanied by a blunted and delayed rise in circulating human platelets. These studies demonstrate that the lung entraps and processes both circulating Mks and released large cytoplasmic fragments consistent with a prior lung/heart murine study and support a pulmonary-centric “catch-and-release” model of thrombopoiesis. Thrombopoiesis is a drawn-out process with the majority of cytoplasmic processing of Mks occurring in the pulmonary bed.

**Key Points:**

- Infused *in vitro*-differentiated megakaryocytes synchronously release cytoplasmic fragments selectively in the pulmonary bed.
- Large, released megakaryocyte fragments recycle to the lungs, undergo further fission, terminally forming platelets.

## Introduction

Mammals have circulating, small anucleate cytoplasmic fragments termed platelets, rather than larger, nucleated thrombocytes^1^, decreasing blood viscosity and increasing hemostatic efficiency in a high-velocity, high-pressure vasculature. Platelets arise from the cytoplasm of polyploid, large megakaryocytes (Mks) found mostly in the marrow^2,3^. The release of platelets from Mks is termed thrombopoiesis and is central in maintaining a physiologic platelet count. A better understanding of thrombopoiesis may improve treatment for clinical states associated with too few or too many platelets and in production of platelets from *in vitro*-differentiated Mks^4^.

One would expect that after studying Mks and platelets for over 100 years^5^, that where thrombopoiesis occurs would be settled. Early *in vitro* megakaryopoiesis videos showed Mk proplatelet extensions greatly influenced the concept of thrombopoiesis^6^. These studies were followed by *in vivo* two-photon microscopy studies in mice (m), suggesting a more complicated series of events where mMks migrate from the marrow into venous sinuses, releasing large cytoplasmic fragments and occasional whole Mks^7–9^. *In situ* studies of the lungs of C57BL/6/PF4-Cre/Rosa26-LSL-tdTomato/mTmG/nTnG/BoyJ (PF4 Cre-mTmG) mice showed large mMks being immobilized in the lungs, releasing large cytoplasmic fragments^10^. The contribution of this observation to overall thrombopoiesis has been debated^11^. A study of recycled mMks in a lung/heart-*in situ* model supported that the lungs entrap Mks, eventually thousands of platelet after recycling multiple times^12^.

Previously, we showed that intravenously (IV)-infused mouse or human (h) Mks into mice resulted in immediate pulmonary entrapment and a peak platelet release 1.5-4 hours post-infusion13,14. These platelets were physically and functionally similar to donor-derived- (dd)-platelets. Using pulmonary *in situ* microscopy, we now confirm the prior observation that endogenous mMks are spontaneously immobilized in the lungs of PF4 Cre-mTmG mice, releasing large cytoplasmic fragments^10^. We then extended these *in situ* studies using IV-infused, *in vitro*-differentiated mMks and hMks into immunocompromised NOD-SCID gamma (NSG) mice. Our findings with infused mMks were consistent with those of the lung/heart-*in situ* studies^12^. In contrast, infused, mature hMks were enucleated upon their first entrapment. Similar studies were done with other hMks: lipopolysaccharide (LPS)-induced inflammatory hMks (inflam-hMks)^15^, umbilical cord-blood-derived (UCB)-hMks, and immortalized Mk progenitor cell (imMKCL)-derived hMks^16^. Exposure of hMks and other cells with blebbistatin, a non-muscle myosin II ATPase inhibitor^17^, and triacsin C, an inhibitor of long-chain fatty acyl-CoA synthetase^18^, support that membrane stiffness is important for proplatelet formation in the pulmonary bed. To address whether the pulmonary bed contributed uniquely to thrombopoiesis or was just the first bed encountered, we compare infusing hMks IV to intraarterial (IA). We propose a multiple-step, pulmonocentric model of “catch-and-release” thrombopoiesis model and discuss its implications.

## Materials and Methods

### In situ-microscopy studies

#### Endogenous mMks studies

*In situ*, real-time-microscopy studies of endogenous mMks entrapment in the lung and the release and recycling of cytoplasmic fragments were done using PF4 Cre-mTmG mice in whom endogenous mMks and expressed green fluorescent protein (GFP)^10^. Mice were anaesthetized with Nembutal. A tracheal cannula was inserted and attached to a MiniVent ventilator (Harvard Apparatus). Mice were ventilated at 130-140 breaths per minute and 10 µl of air/gram of mice with a positive-end expiratory pressure of 2 cm H_2_O. The mice had the surface of the left lung exposed. A thoracic window was applied and 25 mm Hg of suction was applied using a Vacuum Regulator (Amvex) to immobilize the lung. The confocal microscope 50X objective was lowered onto the thoracic window.

#### Infused cell studies

Either dd-platelets or *in vitro-*grown mMks and hMks or bone marrow mononuclear cells (BMMC, Lonza) or mesenchymal stem cells (MSC, Lonza) were infused. Prior to these infusions, cytoplasm was stained with Calcein-AM (C1430, ThermoFisher)^19^ and nuclei with either Hoechst 33342 (H1399, ThermoFisher) or Vybrant DyeCycle Ruby (V10273, ThermoFisher)^19,20^. Mice were also pre-infused with Dextran, Texas Red (D1830, ThermoFisher) to delineate lung vasculature.

8-10X10^6^ cells in 200 µl of culture media were infused into 8-to-12-week-old recipient NSG mice via the jugular vein over 20-30 seconds. Images were analyzed using ImageJ (ver. 0.5.7, NIH) and Imaris (Oxford Instruments). ImageJ was used for the quantification of size (area), shape (aspect ratio), and count of particles, and the results were visualized using R Studio ver.4.2.2. Intrapulmonary released nuclei of infused Mks were analyzed by tracking with Imaris.

Cells were sized by flow cytometry using 4µ to 25µ beads (Invitrogen). We tested the importance of cell fluidity by altering the membrane lipid composition using either triacsin C (3µM), a long-chain acyl-CoA synthetase (ACSL) inhibitor used as described and added on Day 11 of differentiation^21^ or by altering the cytoskeleton with blebbistatin (20µM), a myosin II inhibitor, added on Day 11 of differentiation^22^.

### Analysis of entrapped Mk size and elongation asymmetry

Entrapped Mk size was determined by its number of pixels. Elongation asymmetry was measured by the ratio of width/height. Snapshots videos at specific timepoints after Mk infusion were analyzed from 3 separate studies. A polynomial regression line was calculated by program R (ver. 4.2.2.) that indicates how the dependent variable changes as the independent variable changes^23^. The 95% confidence interval around the regression line was calculated by ggplot2^24^.

### Circulating Mks, cytoplasmic fragments and platelets

Circulating Mks, cytoplasmic fragments and platelets were studied in heparinized (1Unit/20-gram mouse). Isolated mouse and human platelets or Mks were stained with either FITC-labeled rabbit anti-mCD41 (BD Pharmingen) or APC-labeled rabbit anti-hCD41 (BD Pharmingen) antibody prior to infusion. Infused Mks were also pre-labeled with Hoechst-33342 for documenting nuclear presence. Post-infusion of these cells, 30 µl of retro-orbital blood was withdrawn from the mice into EDTA-Micro Capillary Blood Collection (125 µl, Ram Scientific) up to 20 hours post-infusion. Forward scatter (FSC) on flow cytometry was done to measure size of nucleated and anucleated cells and cell fragments.

### IV vs. IA Mk infusion studies

To determine whether IV and IA infused hMks had similar fates, we infused 3-5X10^6^ hMks into NSG mice via a jugular vein or the left carotid artery. Organs were harvested 10 minutes or 4 hours post-infusion and fixed in 10% neutral-buffered formalin (24895887, Sigma-Aldrich). Immunohistologic studies were done staining with anti-human nucleoli antibody (ab190710, abcam) and counterstained with hematoxylin and eosin. Human platelet counts were concurrently measured by flow cytometry.

See the **Supplement Materials and Methods** for a description of the preparation of platelets, Mks and other cells, Institutional approval of human and animal studies, and statistical analysis.

## Results

### Studies of endogenous and infused mMks pulmonary entrapment

We affirm the pulmonary *in situ* microscopy studies done in PF4 Cre-mTmG mice^10^. We observed a mean ± 1 standard deviation (SD) of 2.1 ± 1.0 endogenous mMks entrapped per 50X magnification field per hour (Figure 1A, and Videos 1 and 2). Entrapped mMks formed cytoplasmic extensions with near-complete cytoplasmic release by 40 minutes as previously reported^10^.

**Figure 1.**
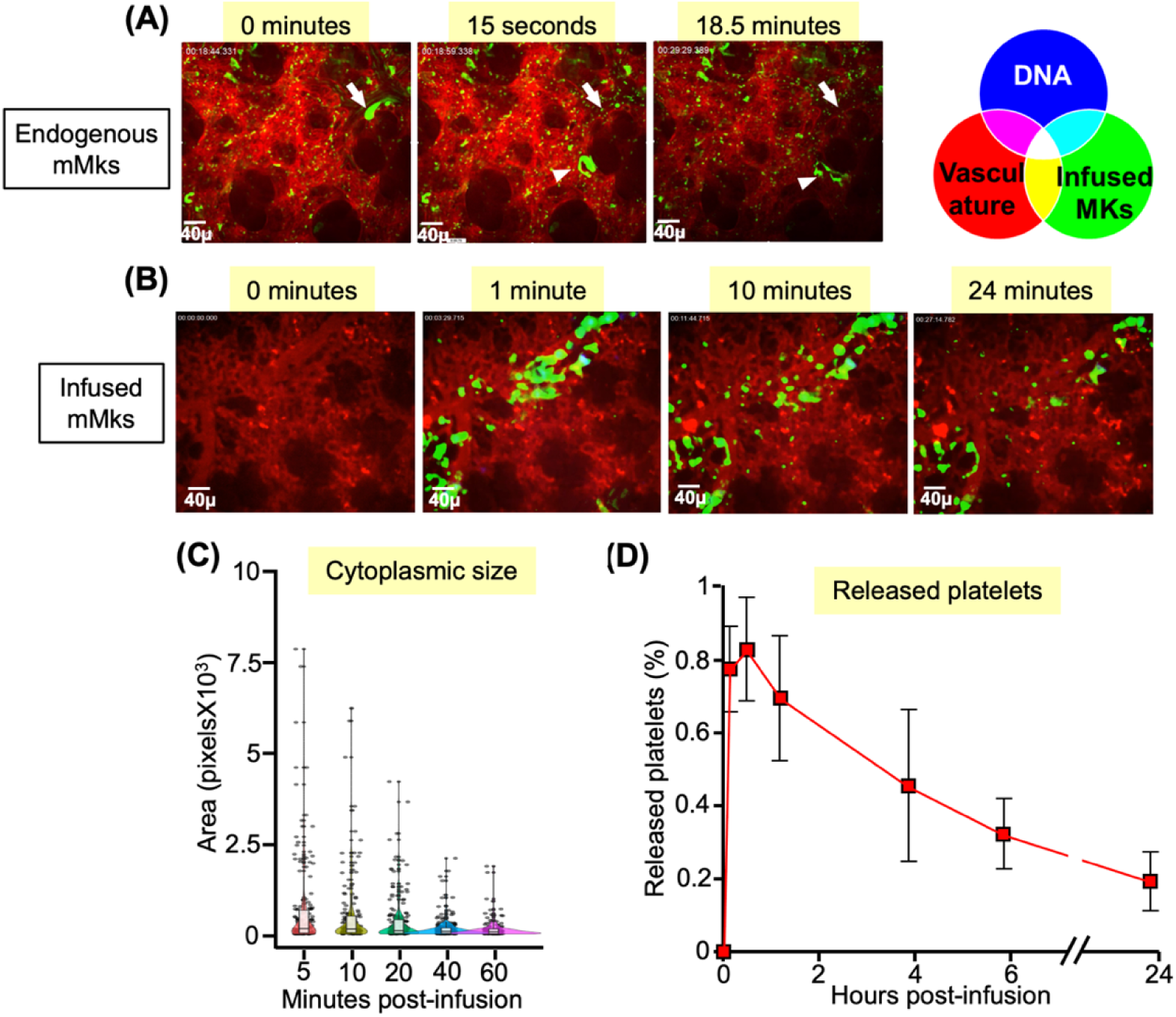
Analysis of endogenous mMks and infused, *in vitro*-grown mMks immobilized in the pulmonary vasculature. (**A**) Representative snapshots beginning at the time of initial appearance of an endogenous mMk shown in Video 1 done in PF4 Cre-mTmG mice until the time of cytoplasmic dispersion. White arrow is the point-of-entry of the mMk, and white arrowhead is the point of cytoplasmic dispersion. Size-bar is included here and in all lung snapshots in this and subsequent figures. Also, see Video 2. N=3 separate mice were studied by video. (**B**). Same as in (A), but snapshots were from Video 3 after infusion of 8X10^6^ *in vitro*-grown WT, Calcein-AM- and Vybrant DyeCycle Ruby-labeled nuclei of mMks infused via a jugular vein into WT mice. Also, see Figure S1 for schematic of methodology and Video 4. N=3 separate mice studied by video after infused mMks. (**C**) Individual cell size was determined by area (i.e., number of pixels) at timepoints 5-60 minutes post infused mMks. The width of each colored shaded area indicates the number of fragments of that particular size at that timepoint. For each timepoint, the mean is shown as well as gray bar indicating ± 1 standard deviation (SD). Determined from three separate infusion. (**D**) Circulating mouse platelets within the FSC window for mice platelets that were derived from infused mMks relative to the total circulating mice platelet population. Mean ± 1 SD shown. N=3 separate animals studied.

We had shown that IV-infusion of *in vitro*-differentiated mature mMks resulted in the release of platelets from entrapped Mks in the lungs with peak platelet counts at 1.5-4 hours^13^. We now infused *in vitro*-grown wildtype (WT) mMks into recipient NSG mice (Figure Supplement (S) 1) and observed that these mMks were entrapped in the pulmonary bed (Figure 1B, and Videos 3 and 4), forming 2.3 ± 0.8 cytoplasmic extensions per mMk that were ∼45 µm in length (Table 1). Peak cytoplasmic extension occurred 10 minutes post-infusion (Figure S2A), and being complete at 40-60 minutes (Figure 1C). The peak rise of released platelets was at 90-minutes (Figure 1D) and disappeared with a half-life of ∼4 hours, consistent with infused murine dd-platelets^13^.

**Table 1.** Characteristics of entrapped Mks. Data represent means ± SD. N=3 videos studied/arm. *p, .05, **p< .005 compared to mature hMks. ╇*p*< .05, ╇╇*p*< .005 compared to mMks.

|  | Endogenous mMks | Infused mature hMks | Infused immature hMks | Infused inflam-hMks | Infused UCB-hMks | Infused imMKCL-hMks |
| --- | --- | --- | --- | --- | --- | --- |
| Cytoplasmic fragments per Mk | 2.3 ± 0.8 | 2.7 ± 0.3 | 0.6 ± 0.2**† | 1.9 ± 0.6 | 1.7 ± 0.3* | 1.1 ± 0.2*† |
| Cytoplasmic fragments (μm) | 44.7 ± 5.1 | 51.6 ± 10.7 | 18.8 ± 1.9*† | 40.0 ± 10.0 | 34.3 ± 7.0 | 24.3 ± 1.8*† |
| % of Mks releasing fragments | 88.7 ± 4.5 | 93.3 ± 3.8 | 32.3 ± 5.0**†† | 70.3 ± 12.7 | 71.3 ± 6.5*† | 50.0 ± 9.8*† |

### Studies of thrombopoiesis in the lungs from infused CD34^+^-derived hMks

We had shown that infused, *in vitro*-differentiated Day 12, hMks became immobilized in the lungs and released platelets with similar kinetics as infused mMks^14^*. In situ* microscopy studies were now done after IV-infusion of Day 12-14 days hMks differentiated from CD34^+^ cells (Figure S1). Comparable to infused mMks, the hMks were entrapped in the pulmonary vasculature, synchronously extended a similar number and length cytoplasmic extensions (Figure S2), and completed cytoplasmic shedding in <40 minutes (Figures 2A and 2B, Videos 5 and 6, and Table 1). Peak circulating human platelet count was also comparable (Figure 2D).

**Figure 2.**
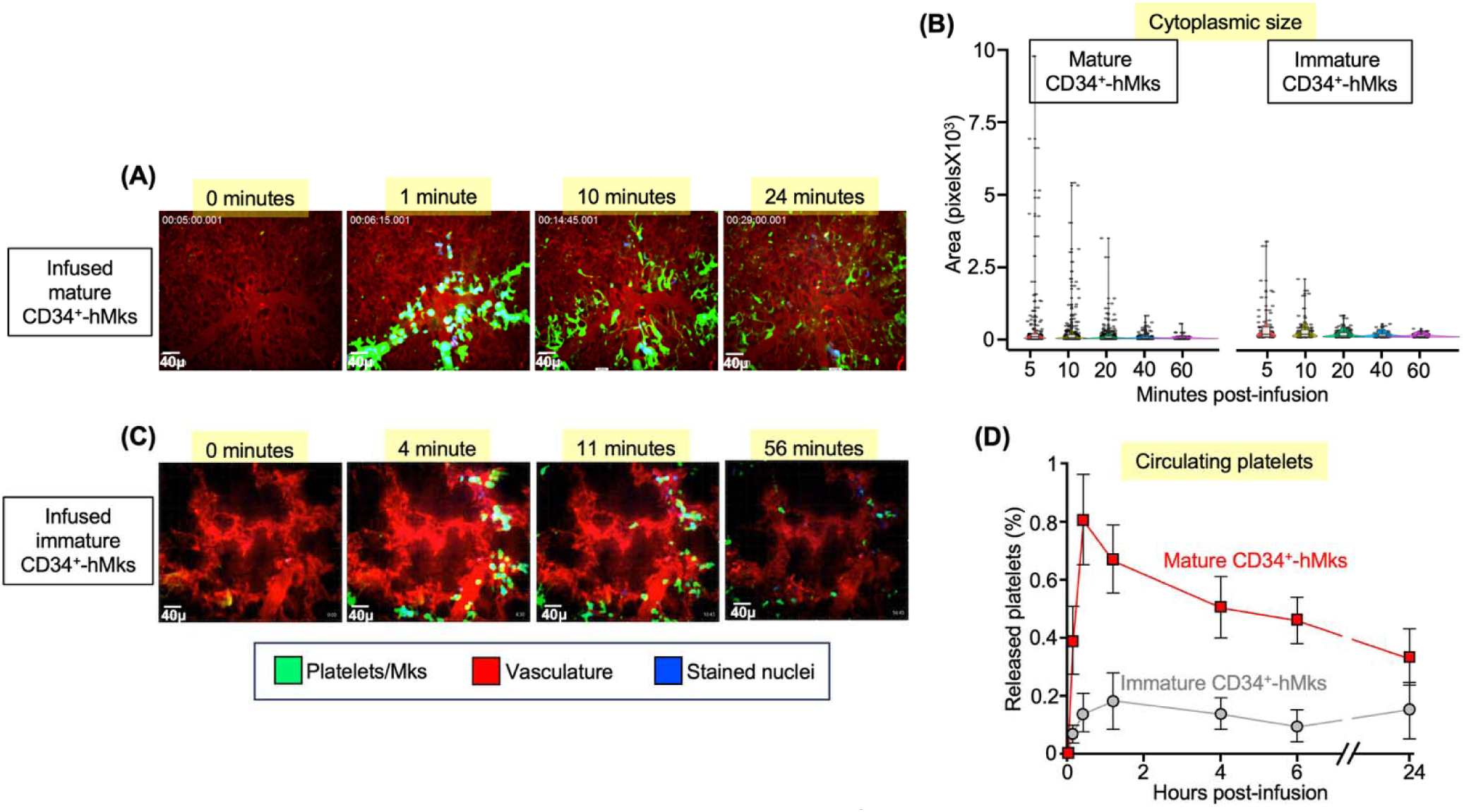
Studies of infused mature and immature CD34^+^-hMks into mice. (A) Representative snapshots of Video 5 done after infusing mature (Day 12) *in vitro*-differentiated CD34^+^ cells to hMks into NSG mice as in Figure S1. See also Video 6 of a similar study. Analysis in (B) and Table 1 is based on N=3 videos of mature hMk infusions. (**B**) Cytoplasmic size in pixels. Same as in Figure 1B, but for infused mature and immature hMks. (**C**) Same as in (A), but from Video 7 after infusing immature (Day 7) *in vitro*-differentiated hMks. See also Video 8. N=3 videos of mature hMk infusions were studied for analysis in (**B**) and Table 1. (**D**) Same as in Figure 1D, but for the released human platelets labeled by anti-hCD41 antibody within the FSC window for dd-platelets.

We also infused immature, Day 7 hMks. Immature hMks were smaller, and had lower surface CD42a and ploidy (Figure S3). Immature hMks, immobilized in the lungs, had limited cytoplasmic shedding (Figure 2C, and Videos 7 and 8). Cytoplasmic extensions were fewer and shorter (Table 1). Extension formation was limited and peaked early post-infusion (Figure S2C) with significantly less extension than mature hMks (Figures 2 and S4, and Table 1). As would be expected, few platelets were released, and the peak human platelet count was delayed (Figure 2D).

### Recirculating Mks and cytoplasmic fragments

The *in-situ* microscopy videos of the steady-state PF4 Cre-mTmG mice pulmonary bed Videos 1 and 2 differed in an important way from mMk Videos 3 and 4 and hMk Videos 5 and 6: There was a heavy background of circulating platelets in addition to the sudden appearance of an occasional large mMks (≥20µ), but in addition, there was a steady level of large (≥12µ to <20µ) and small (≥5µ to <12µ) anucleate cytoplasmic fragments (Figure S5). These fragments can be seen undergoing further division, consistent with Mks undergoing repeat cycles of “catch and release” of larger cytoplasmic fragments in the lungs, undergoing further divisions.

We also stained the Mk nuclei. After infusing mMks or hMks, their stained nuclei were noted in the lungs (Figure 3), but differed in that mMks only had a small initial drop of free nuclei of ∼6% in the first 5 minutes post-peak, while hMks underwent a brisker initial release of their nuclei with an ∼35% drop in nuclei over the first 5 minutes (Figure 3A *vs*. 3B). Either free nuclei – devoid of cytoplasm – or nucleated Mks disappeared systemically (see Figure 3C and Video 9). After 15-20 minutes, few mature mMk or hMk nuclei were present in the lungs. Nuclear outcome differed when immature hMks were infused: there was a more blunted disappearance of Mk nuclei (Figure 3B). This slower drop in free nuclei after mMks and immature hMks infusions may, in part, be explained by longer pulmonary retention and/or by Mk recirculation as seen in Figure 4. The final fate of the circulating free Mk nuclei needs further study.

**Figure 3.**
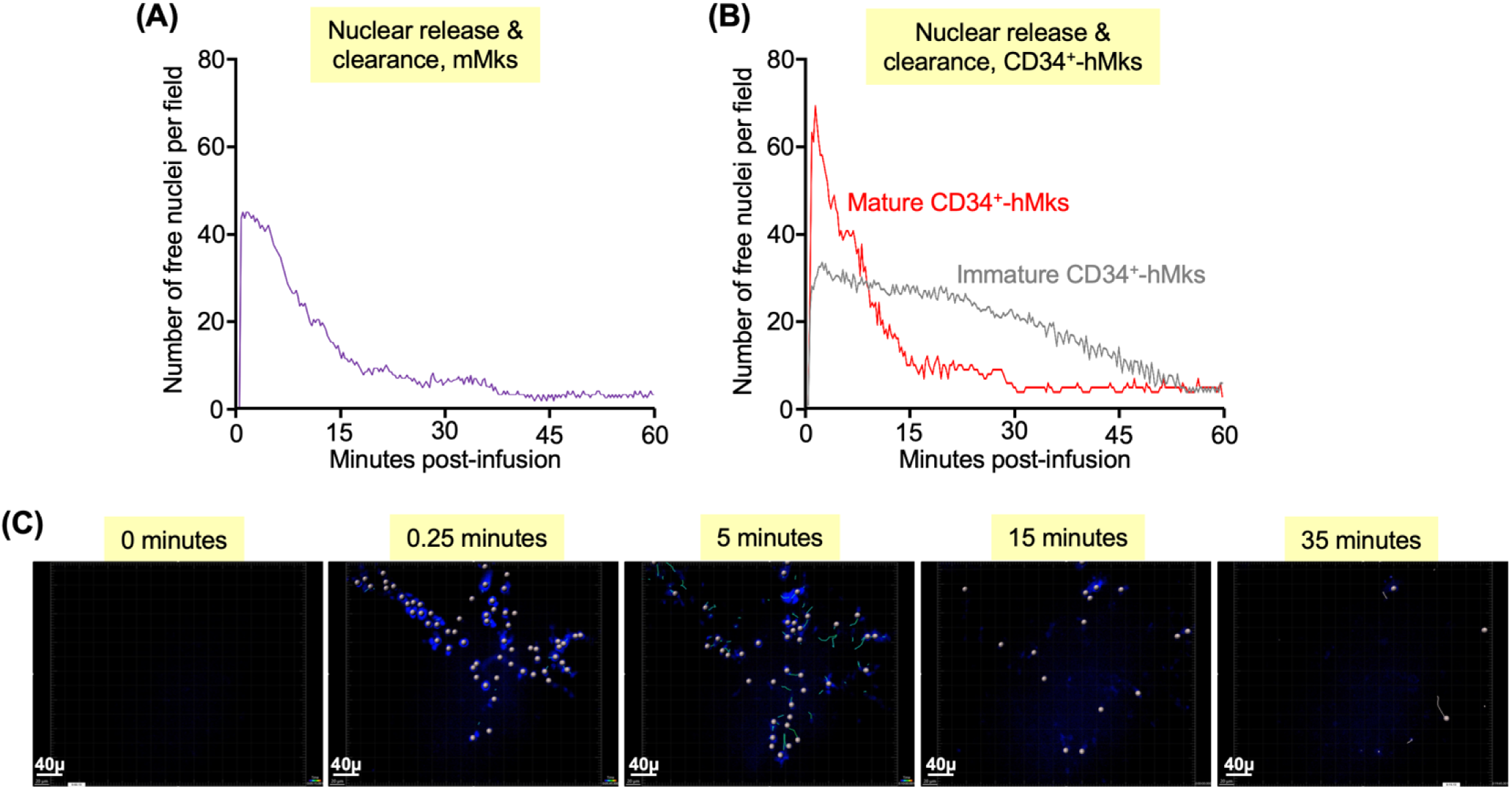
Intrapulmonary released nuclei after infusion of Mks. (**A**) and (**B**) are studies of all stained nuclei from the infused Mks using Vybrant DyeCycle Ruby. In each case, the mean of free nuclei is shown from three separate infusions. (**C**) Snapshots from a study of nuclei followed over time beginning at the time of hMk infusion. A small dot has been placed over each nuclei and its velocity and direction in that snapshot is indicated by a green line. See also Video 9, which shows movement of shed nuclei in blue over this one-hour study post hMk infusion.

**Figure 4.**
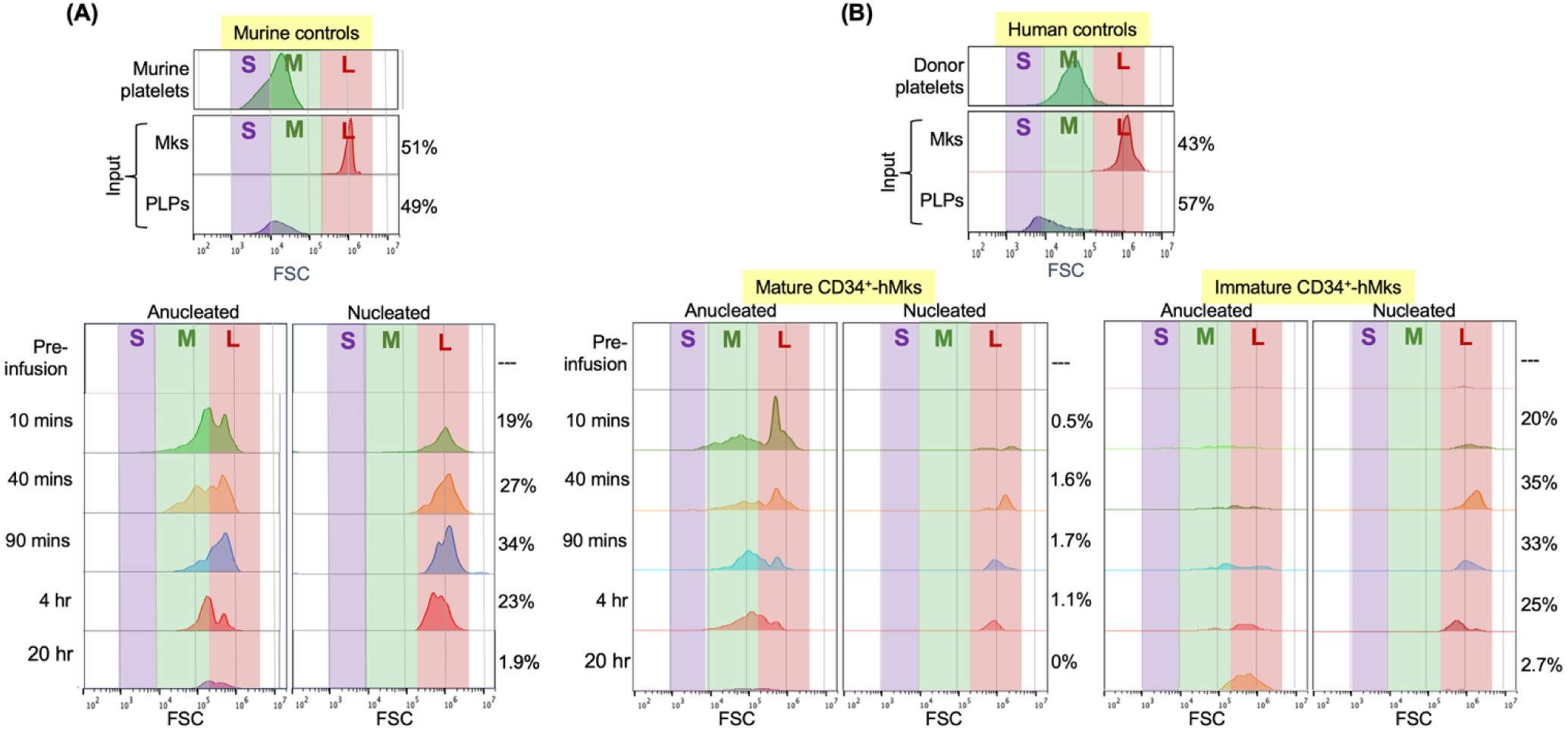
Circulating Mks, cytoplasmic fragments and PLPs post infusion. Peripheral blood studies of *in vitro*-differentiated mature mMks and hMks, and immature hMks sized by FSC. The presence of nuclei was determined by Hoechst blue-staining and expressed as percent of total particles on the ordinate and FSC on the abscissa. All experiments are the mean of three separate studies. (**A**) The top graphs show size controls of isolated murine platelets, and the infusate which is a mix of nucleated mMks and anucleated PLPs. The FSC was arbitrarily divided into three size ranges: S=small size that included mostly PLP-sized particles. M=medium size that included mostly platelets. L=large size that included the nucleated Mks. The same three size ranges are shown at the bottom graphs which shows temporal changes in nucleated and anucleated cells at various times post-infusion. The ordinate represents cell counts and ranges from 0 to 100. (**B**) Similar studies to (A), but for human studies. The top graphs are for dd-platelets and the harvested hMks just prior to its infusion. The ordinate represents cell counts and ranges from 0 to 4856. At the bottom, the left is after infusion of mature hMks and the right after immature hMks. The ordinate again represents cell counts and ranges from 0 to 824 for mature Mks and 0 to 139 for immature Mks.

The lung/heart-*in situ* study suggested that mMks recycled multiple times^12^. To confirm and extend those findings, we examined recycling nucleated Mks and anucleate, cytoplasmic fragments in the peripheral blood of mice after infusion of double-labeled mMks and hMks, and immature hMks. The presence of a nucleus distinguished Mks from large cytoplasmic fragments (Figure S6). As controls, we determined size-distribution of mice and human dd-platelets, and size distribution and nucleation status of the *in vitro*-grown mMks and hMks preparations (top of Figures 4A and 4B, respectively). Both murine and human dd-platelet size followed a bell-shaped curve, and the Mk preparations contained roughly half large, nucleated Mks and half small, anucleated fragments whose size-range overlapped with dd-platelets, but peaked at a smaller size. These particles were termed platelet-like-particles (PLPs) to distinguish them from dd-platelets.

After infusion of the mMks, there was a large pool of circulating nucleated mMks detectable at four hours, but gone the next day (Figure 4A, bottom, right). mMks became smaller with time, consistent with the lung/heart-*in situ* study^12^. A pool of anucleated, circulating large cytoplasmic fragments was prominent at 10 minutes, but replaced by a smaller fragments – still larger than murine platelets – by 4 hours (Figure 4A, bottom, left). The PLPs in the mMk infusate were quickly cleared by 40 minutes.

Similar studies were done for mature and immature hMks (Figure 4B). In contrast to mMks, after infusing hMks, the pool of recycling nucleated cells was minor; however, the few recycling hMks still showed a steady decline in size, suggesting that they were shedding cytoplasm over time. The largest pool at the earliest timepoint was an anucleate, large-cytoplasm pool. This compartment steadily decreased in mean size and in prominence over time (Figure 4B, bottom left). A platelet-size pool was present at the earliest timepoint and became the prominent pool by 4 hours. The smaller-size PLP pool is only present at 10 minutes, consistent with PLPs being preactivated and rapidly cleared^14^. Immature hMks behaved more like infused mMks in that there was a large fraction of recirculating nucleated Mks (Figure 4B, right). Immature hMKs released a modest large-cytoplasm fragment pool and a platelet-sized pool.

### Thrombopoiesis in the lungs from infused inflam-hMks

Mks have been described during hematopoiesis, which have a more immune protein signature and function^25^. The capacity of these inflam-hMks to release platelets has been understudied. We induced inflam-hMks from CD34^+^-HSPCs using LPS^26^ with increased surface levels of major histocompatible complex (MHC) II used to identify these cells^26,27^ (Figure S7). Inflam-hMks retained the ability to be immobilized in the lungs, and extend and release large cytoplasmic fragments (Figures 5A and 5B *vs*. 2A and 2B, Table 1, and Videos 10 and 11). Inflam-hMks formed comparable numbers and length of cytoplasmic extensions as adult hMks and in the same timeframe (Figure S2D and Table 1). Nuclear release and clearance and platelet release were normal (Figure S8 and 5C, respectively). Thus, our data would suggest that inflam-hMks undergo efficient thrombopoiesis and contribute to the peripheral platelet count.

**Figure 5.**
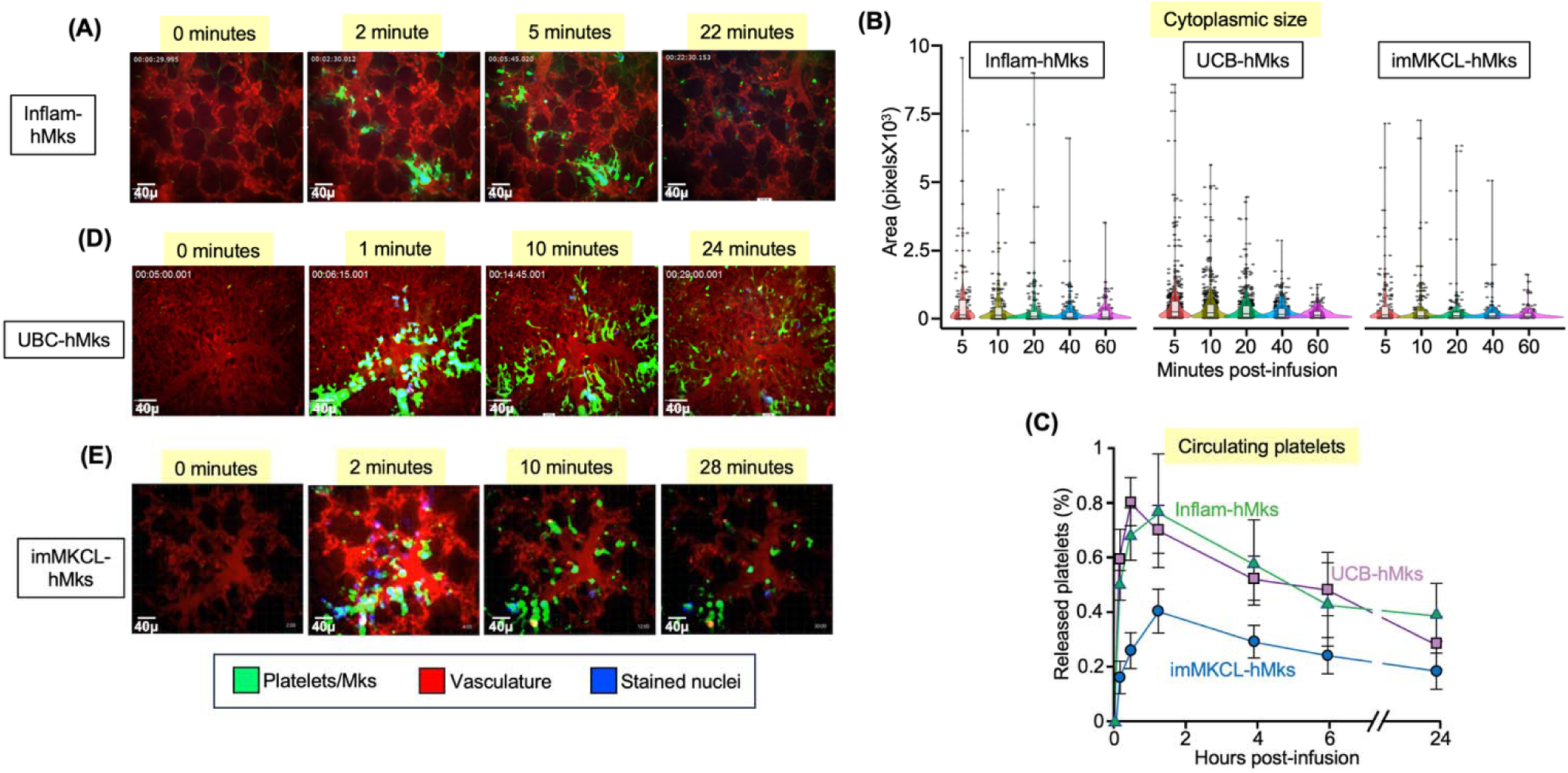
Studies of hMks derived under other clinical conditions. (**A**)-(**E**) Studies similar to studies in Figure 3, but for inflam-, UCB- and imMKCL-hMks. (A), (D) and (E) are video snapshots from Videos 10, 12, and 14, respectively, after infusion of the *in-vitro* differentiated hMks. For the various infused hMks studied, (B) is cytoplasmic size over time as in Figure 1C, and (C) is the mean ± 1 SD for the number of platelets released relative to the murine platelet count as in Figure 2D. N=3 separate animals studied.

### Thrombopoiesis in the lungs from UCB-hMks and imMKCL-hMks

Fetal circulation largely bypasses the lungs, but this changes upon birth^28^. We asked whether infused UCB-hMks would efficiently undergo pulmonary thrombopoiesis. While smaller than adult hMks^29^, infused UCB-hMks were efficiently immobilized in the lung and synchronously shed their cytoplasm in <40 minutes (Figures 5B and 5D *vs*. 2A and 2B, and Videos 12 and 13). UCB-hMks formed comparably long cytoplasmic extensions as adult hMks though somewhat later (Figure S2E and Table 1). Nuclear release and clearance were similar to adult hMKs (Figure S8). Circulating platelet levels after UBC-hMk infusion were comparable to that seen with infused adult hMks (Figure 5C *vs*. 2D).

Induced-pluripotent stem cells (IPSCs) can be differentiated into Mks, but these Mks are thought to have an embryonic signature^30^. The cell line imMKCLs is an iPSC line partially differentiated towards hMks that completes differentiation upon doxycycline withdrawal^31^. Platelets from similar cells were reported in a first-to-human infusion trial^32^. Like immature, adult hMks, imMKCL-hMks were small and formed few extensions (Figure 5E *vs*. 2A, and Videos 14 and 15). Cytoplasmic extension was delayed (Figures S2E and 5B), and their size blunted (Table 1). Nuclear clearance and platelet release were reduced (Figures S8 and 5C, respectively).

### Comparative thrombopoiesis studies of IV- to IA-infused mature hMks

Is the pulmonary vascular bed unique in entrapping Mks and releasing cytoplasmic fragments or is it simply the first vascular bed encountered by released Mks and/or large cytoplasmic fragments from the bone marrow and other Mk-producing organs^33^? Would IA-infused hMks release cytoplasmic fragments and contribute to circulating platelets? To address these questions, we infused mature hMks IA via the left carotid artery, occluding the cannulated vessel so that the infused Mks retrogradely dispersed, or IV via the tail vein, and followed both the release of circulating human platelets and the presence of immobilized hMks in various organs (see schematic in Figure S9). The rise in platelet counts post-IV infusion (Figure 6A) was similar to what we reported^14^ and in Figure 2D. In contrast, post-IA infusion, there were almost no detectable circulating human platelets initially. Low levels were detectable at 4 hours. Post-IA infusion, there were almost no Mks in the lungs at 10 minutes, but were clearly notable at 4 hours (Figure 6B). Aside from an occasional Mks in the liver at 10 minutes, no Mks were seen in the other tissues examined post-IA infusion (Figures 6C, 6D, and S10).

**Figure 6.**
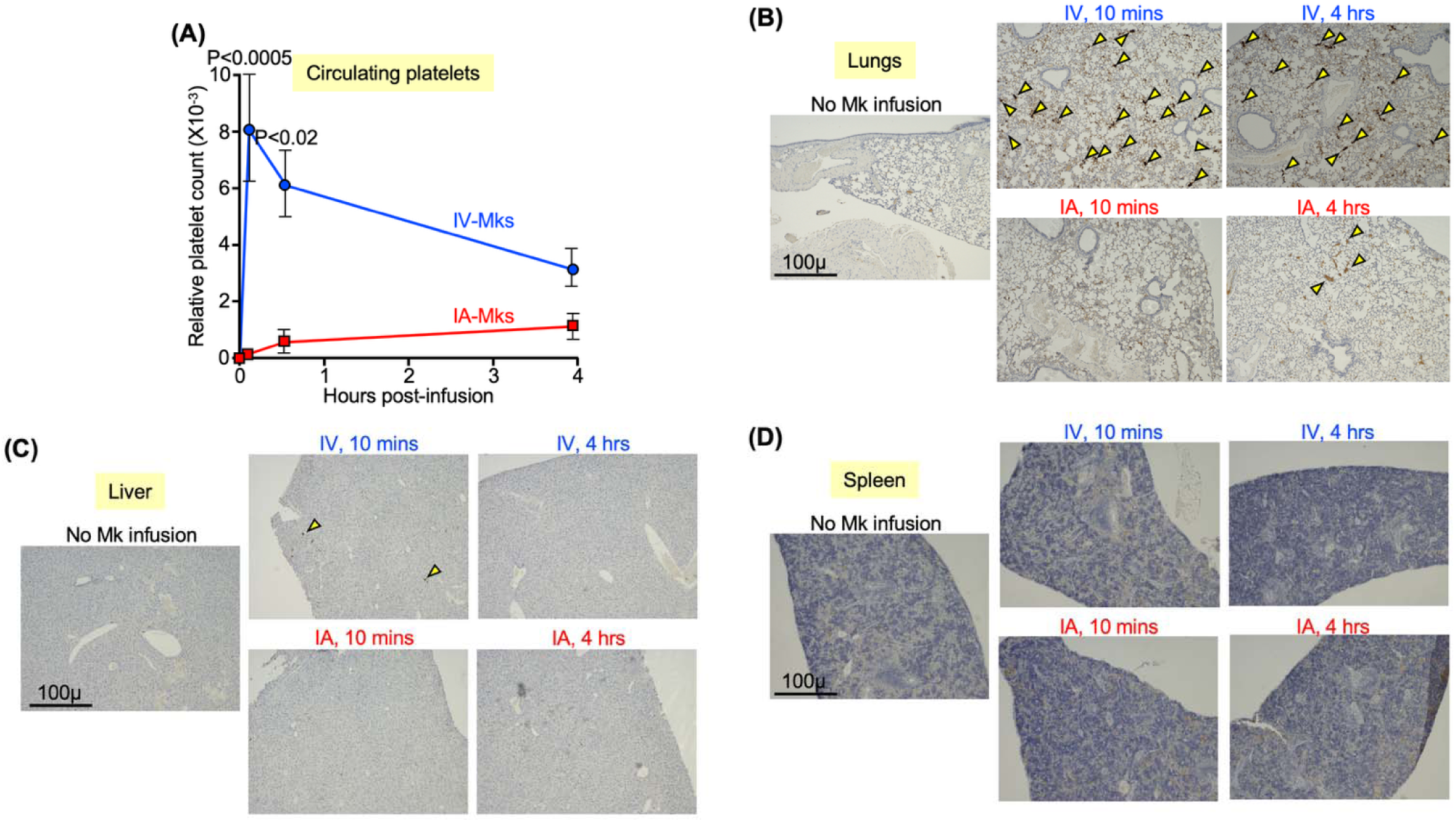
Consequences of IV *vs*. IA infused hMks into NSG mice. To examine the importance of the pulmonary vasculature in thrombopoiesis, the same number of *in vitro*-differentiated mature hMks were infused into NSG mice IV or IA. (**A**) Changes in the number of circulating human platelets over time was followed. N=5 distinct mice per arm studied. Mean ± 1 SD is shown over the first 4 hours post-infusion. Significant differences at specific timepoints were determined by two-way Student t test. (**B**)-(**D**) Representative immunohistology stained with an anti-human nuclear antibody of the lungs, liver and spleen, respectively, from control NSG mice not receiving hMks or 10 minutes or 4 hours post-infused hMks. Yellow arrowheads indicate observed hMks entrapped in specific organs. Size bars are included in (B)-(D).

### Mk properties and pulmonary thrombopoiesis

The mechanico-biologic processes that lead to intrapulmonary thrombopoiesis include properties intrinsic to the immobilized Mks, physical organization of and receptors lining the pulmonary vasculature, gaseous exchange occurring in the lung, altering oxygen and CO_2_ levels and pH, and inhalation and expiration that alter the cross-sectional diameter of pulmonary vessels^12^. We examined physical characteristics of Mks to see whether they contribute to intrapulmonary thrombopoiesis including whether the size of the Mks contribute to the process by examining BMMCs that are smaller than UCB-hMks, and MSCs that are larger than mature CD34^+^-hMks (Figure S11A). BMMCs were not entrapped in the lungs (Figure S11B, and Videos 16 and 17), while MSCs were immobilized in larger arterioles and remained spherical (Figure 7A, and Videos 18 and 19). Prior studies have shown that blebbistatin, enhances *in vitro* PLP formation from Mks^34^. Treatment with blebbistatin^17^ led to the MSCs becoming embedded in the smaller pulmonary vessels similar to CD34^+^-hMks and were no longer spherical, having cytoplasmic projections (Figure 7A and Videos 20 and 21) had circulating PLP-like fragments not seen with untreated MSCs (Figure 7B). Blebbistatin-exposed hMks had enhanced proplatelet projections (Figures 7A and 7C and Table 2), but imMKCL cells treated with blebbistatin did not have enhanced proplatelet extensions and only a modest increase in platelet release compared to untreated imMKCLs (Figures 7A and 7D and Table 2).

**Figure 7.**
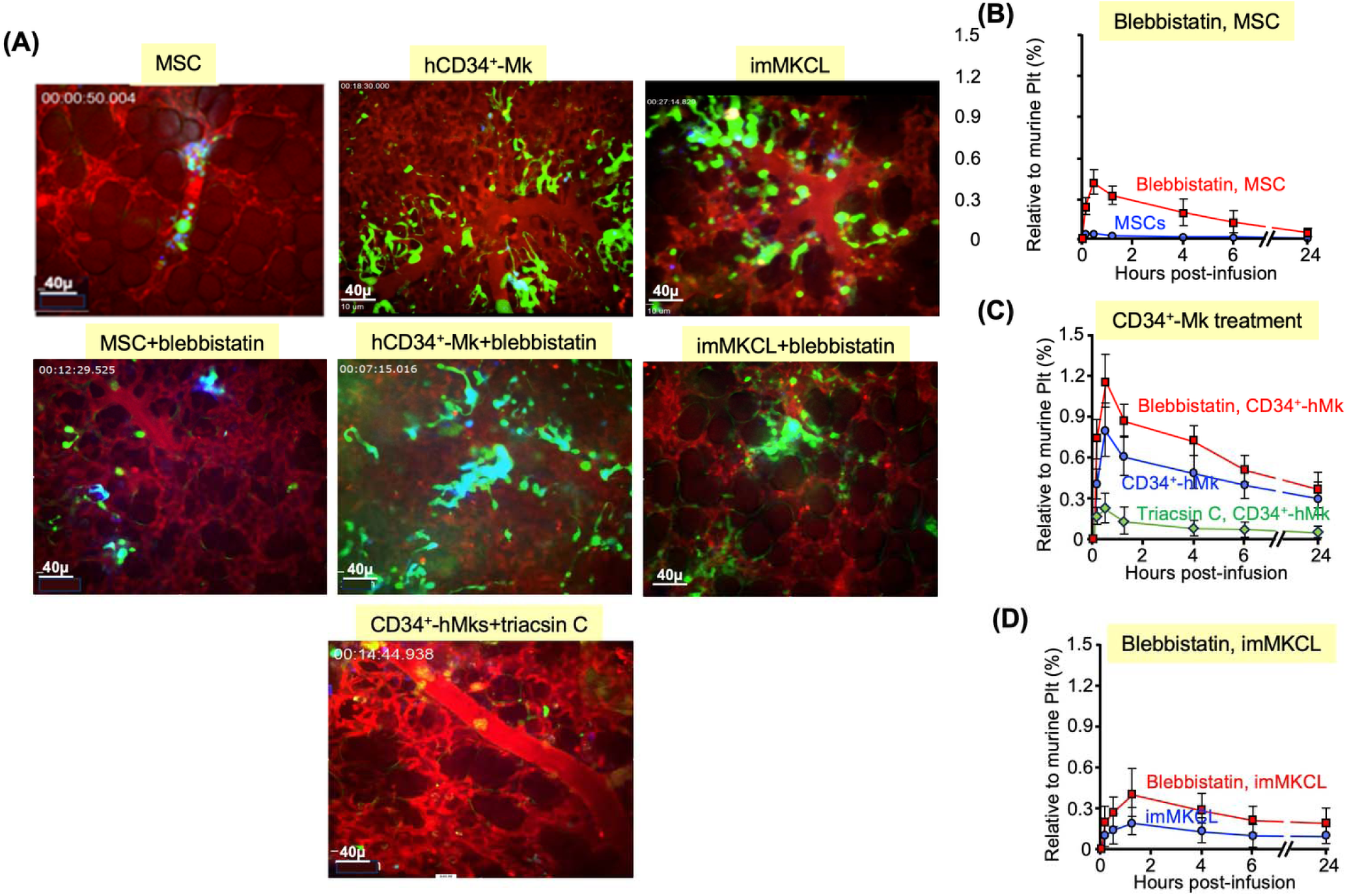
Effects of blebbistatin and triacsin C on thrombopoiesis. (**A**) Representative snapshots of infused MSCs, CD34^+^-hMks and imMKCL-hMks and entrapped in the pulmonary bed without treatment (top), with blebbistatin- (middle) or triacsin C-treatment (bottom). See Videos 16 through 23. (**B**)-(**D**) PLPs release after infusing a similar number of cells untreated or treated as described. Mean ± 1 SD over the 24 hours following cell infusion. N=3 studies per arm. (B) MSCs that had or had not been pretreated with blebbistatin. (**C**) Same as (B), but for hCD34^+^-hMks untreated or pre-treated with either blebbistatin or triacsin C. (**D**) Same as (B), but for imMKCL pretreated or not with blebbistatin.

**Table 2.** Blebbistatin and triacsin C effected on hMks. . Data represent means ± SD N=3 videos studied/arm. \**p*<.o5, \*\**p*<.005 compare to mature hMks. ns= not significantly different from imMKCL cell not treated with blebbiststin.

|  | Infused mature hMks | Infused mature hMks + blebbistatin | Infused mature hMks + triacsin C | Infused imMKCL-hMks | Infused imMKCL-hMks + blebbistatin |
| --- | --- | --- | --- | --- | --- |
| Cytoplasmic fragments per Mk | 2.7 ± 0.3 | 3.8 ± 0.8 * | 1.6 ± 0.5 * | 1.1 ± 0.2 * | 1.3 ± 0.5<sup>ns</sup> |
| Cytoplasmic fragments (μm) | 51.6 ± 10.7 | 69.7 ± 11.6 * | 25.5 ± 9.9 * | 24.3 ± 1.8 * | 28.4 ± 5.6<sup>ns</sup> |
| % of Mks releasing fragments | 93.3 ± 3.8 | 97.5 ± 7.4 | 55.2 ± 4.9 ** | 50.0 ± 9.8 * | 54.2 ± 7.9<sup>ns</sup> |

Recently, studies have shown that altered Mk membrane fatty-acid composition can affect thrombopoiesis^35^. In particular, triacsin C inhibited thrombopoiesis. We examined triacsin C^18^-treated hMks immobilization in the pulmonary bed and found that the Mks were now immobilized in the larger arteriole and appeared spherical, similar to entrapped, untreated MSCs (Figure 7A and Videos 22 and 23). These hMks also showed markedly reduced platelet-release (Figure 7C).

## Discussion

The prevailing model of platelet formation in adult mammals is that Mks predominantly differentiate in the marrow and that as they emerge into the bloodstream, Mks release proplatelets that are rapidly converted into platelets^33^. Videos of *in vitro*-differentiated Mks supported this model with an occasional Mk in culture undergoing proplatelet formation of “beads-on-a-string”^6^. Based on this model, multiple papers relied on PLP release from proplatelets in culture as an indicator of the capacity of *in vitro*-grown Mks to undergo thrombopoiesis in various culture conditions^36–38^ or genetic disorders^39–41^. Additionally, devices have been described to take advantage of this proplatelet formation as a potential source of non-dd-platelets^42,43^.

*In vivo* murine studies suggested an alternative model: Two-photon microscopy studies of the diploic calvarium space did not show beads-on-a-string extensions^7–9^, but rather showed large cytoplasmic fragments being released into the medullar veins from one-to-a-few Mk extensions. Yet most reviews medullar venous beads-on-a-string release^44–46^. That large fragments were typically observed and these could remain attached by a thin cytoplasmic thread to each other and to the trailing nucleated body and that occasional, whole Mks were released was underplayed^7–9^. The interconnected-large, cytoplasmic fragments could break apart or reconstitute into intact Mks prior to reaching the lung. The *in situ* pulmonary microscopy study confirmed that large Mks from the bone marrow can be immobilized in the pulmonary bed^10^. These endogenous mMks then shed their cytoplasm over 30-40 minutes. The contribution of this pulmonary thrombopoiesis in the steady-state has been debated^11^, but the recent recirculated mMk lung/heart-*in situ* study^12^ support that such a recycling pulmonary pathway can led to the release of several thousand platelets per mMk.

The presented infused mMk studies are consistent with the recirculated mMk lung/heart-*in situ* study^12^, but the infused hMk studies differed with respect to enucleation. mMks required multiple passages through the lung to fully enucleate, but hMks enucleate on first passage. The basis for this species difference is unclear. Whether the different patterns of enucleation is an artifact due to incomplete i*n vitro* differentiation of one or the other Mks or due to infusion of xeno- *vs*. species-specific Mks or this difference really exists will need further study. The mMk results were similar to immature hMks in their enucleation pattern. Adult hMks are larger than adult mMks^47^, but immature hMks are closer in size to mMks; however, UCB-hMks that are also small enucleate well^29^. Whether there is an inverse relation between capacity to enucleate and Mk size needs further examination.

Our studies show that *in vitro*-differentiated mMks and hMKs are primed to undergo cytoplasmic shedding synchronously post-infusion. In culture, only an occasional cell sporadically undergoes PLP release. We propose that *in vitro*-PLP release is from Mks undergoing terminal apoptosis that utilizes the proplatelet pathway as Mks grown in a plastic dish are bereft of a natural exit to undergo terminal thrombopoiesis^48^. The PLPs are heterogenous, small fragments that are rapidly cleared when infused, consistent with being preactivated or apoptotic. Such PLPs might be clinically useful as an immediate thrombogenic agent.

The large cytoplasmic fragment and the platelet-sized pools are distinct in size with little overlap. We previously showed that the final, released platelet-sized pool has a similar size distribution and circulating half-life in NSG mice as dd-platelets and are almost as responsive to agonists^14^. We propose that upon reduction to a certain size, cytoplasmic fragments undergo proplatelet formation^49^. This final phase was not studied and may not require the pulmonary bed.

Our studies show for the first time that the pulmonary bed differs from other vascular beds in supporting thrombopoiesis from immobilized Mks and cytoplasmic fragments. In this manuscript we addressed Mk-specific features that may influence thrombopoiesis, focusing on the role of cell size and membrane fluidity on pulmonary thrombopoiesis using pharmacologic alteration of Mks and other cells. Our studies suggest that the size of Mks affects immobilization in the correct pulmonary vessels^49^ and their membrane fluidity regulates subsequent pulmonary thrombopoiesis^35^. Studies of the importance of the physical organization of the pulmonary vasculature^50^, the nature of the pulmonary lining^51^, the exchange of gases and vessel expansion and contraction^52^ would require alternative approaches, perhaps utilizing a “lung-on-a-chip” with flexible membranes allowing gaseous exchange.

While UBC-Mks appear able to undergo pulmonary thrombopoiesis in an adult animal, pulmonary thrombopoiesis in the fetus is likely unimportant as the pulmonary bed is largely bypassed. Likely, an alternative vascular bed substitutes for the lungs. Placentation and thrombopoiesis are both uniquely mammalian features, and the placenta is the functional equivalent of the lungs. Detectable Mks have been noted in the placenta, and based on a comparison of the size and number of Mks in the umbilical arteries *vs*. veins, it had been proposed that fetal thrombopoiesis occurs in the placenta^53^.

The lung being central for thrombopoiesis has clinical relevance. Mks bypassing the lungs has been proposed as a cause of thrombocytopenia in cyanotic-heart disease patients^54^. Individuals on pulmonary-bypass devices also have right-to-left shunting that may contribute to the observed thrombocytopenia. Finally, that thrombocytopenia is not observed in many lung disorders, like emphysema, does not detract from a central thrombopoiesis role for the lung. In these diseases, all circulating Mks and cytoplasmic fragments still pass through the residual pulmonary vasculature^55^.

Another clinical implication of a “catch-and-release” pulmonary thrombopoiesis model relates to developing technologies for *in vitro*-generation of platelets to replace dd-platelets as thrombopoiesis is a complex process involving repeated cytoplasmic divisions. Single-passage sieves are unlikely to be effective. Devices involving serial passage through a channel system may be more effective as proposed in the lung/heart-infusion paper^12^ and to continuous turbulence systems that may allow repeated cytoplasmic division^56^.

In summary, studies of murine and human thrombopoiesis in mice support a complex, pulmonocentric model. Mks and their cytoplasmic fragments appear to undergo a “catch-and-release” strategy in the lungs where they are repeatedly immobilized and undergo repeated fragmentation, eventually releasing platelets. While Mks are initially immobilized in the lung for only half-an-hour, complete thrombopoiesis requires 1.5-4 hours. For hMks, enucleation occurs upon first lung entrapment with the nuclei released downstream. For mMKs, enucleation requires multiple cycles of pulmonary entrapment, while hMks appear to release their cytoplasm on first passage. No other vascular bed carries out this “catch-and-release” process. The described model has important implications in understanding thrombocytopenia in clinical settings involving right-to-left lung bypass and in considering how to generate platelets from *in vitro*-grown Mks to substitute for dd-platelet transfusions.

## Acknowledgments

This work was supported by grants to MP from the NHLBI (R35 HL150698) and KL from the National Research Foundation of Korea (NRF) (RS-2025-00556460) and the Hankuk University of Foreign Studies Research Fund. We thank Dr. Mark Looney at the University of California San Francisco for teaching our group the *in situ* pulmonary microscopy system; Dr. Koji Eto at Kyoto University, Center for iPS Cell Research and Application, Kyoto, Japan, for providing the imMKCL cells; Dr. Alastair Poole from the University of Bristol for helpful discussions; and Dr. Douglas B. Cines from the University of Pennsylvania for reviewing the manuscript.

## Contribution to this manuscript

HK carried out most of the described studies, assisted in design and interpretation of studies and in the preparation of the manuscript. HA performed experiments and taught HK Mk culturing technologies. DJ carried out many of the intra-arterial studies. LR established the *in-situ* microscopy system and provided important insights into data interpretation. KL developed the studies and analysis of the circulating megakaryocytes and cytoplasmic fragments, and assisted in the writing and editing of the manuscript. MP developed the underlying concept and provided overall guidance as well as preparation of the manuscript.

## References

1. Canfield PJ. Comparative cell morphology in the peripheral blood film from exotic and native animals. Aust Vet J. 1998;76(12):793–800.

2. Puhm F, Laroche A, Boilard E. Diversity of Megakaryocytes. Arterioscler Thromb Vasc Biol. 2023;43(11):2088–2098.

3. Guo K, Machlus KR, Camacho V. The many faces of the megakaryocytes and their biological implications. Curr Opin Hematol. 2024;31(1):1–5.

4. Chen SJ, Sugimoto N, Eto K. Ex vivo manufacturing of platelets: beyond the first-in-human clinical trial using autologous iPSC-platelets. Int J Hematol. 2023;117(3):349–355.

5. Weyrich AS, Zimmerman GA. Platelets in lung biology. Annu Rev Physiol. 2013;75:569–91.

6. Italiano JE Jr, Lecine P, Shivdasani RA, Hartwig JH. Blood platelets are assembled principally at the ends of proplatelet processes produced by differentiated megakaryocytes. J Cell Biol. 1999;147(6):1299–1312.

7. Junt T, Schulze H, Chen Z, et al. Dynamic visualization of thrombopoiesis within bone marrow. Science. 2007;317(5845):1767–70.

8. Zhang L, Orban M, Lorenz M, et al. A novel role of sphingosine 1-phosphate receptor S1pr1 in mouse thrombopoiesis. J Exp Med. 2012;209(12):2165–81.

9. Brown E, Carlin LM, Nerlov C, Celso CL, Poole AW. Multiple membrane extrusion sites drive megakaryocyte migration into bone marrow blood vessels. Life Sci Alliance. 2018;1(2):e201800061.

10. Lefrançais E, Ortiz-Muñoz G, Caudrillier A, et al. The lung is a site of platelet biogenesis and a reservoir for haematopoietic progenitors. Nature. 2017;544(7648):105–109.

11. Malara A, Balduini A. Megakaryocytes in the lung: guests or ghosts? Blood. 2024;143(3):192–193.

12. Zhao X, Alibhai D, Walsh TG, et al. Highly efficient platelet generation in lung vasculature reproduced by microfluidics. Nat Commun. 2023;14(1):4026.

13. Fuentes R, Wang Y, Hirsch J, et al. Infusion of mature megakaryocytes into mice yields functional platelets. J Clin Invest. 2010;120(11):3917–3922.

14. Wang Y, Hayes V, Jarocha D, et al. Comparative analysis of human ex vivo-generated platelets vs megakaryocyte-generated platelets in mice: a cautionary tale. Blood. 2015;125(23):3627–3636.

15. Guo L, Shen S, Rowley J, et al. Platelet MHC class I mediates CD8+ T-cell suppression during sepsis. Blood. 2021;138(5):401–416.

16. Nakamura S, Takayama N, Hirata S, et al. Expandable megakaryocyte cell lines enable clinically applicable generation of platelets from human induced pluripotent stem cells. Cell Stem Cell. 2014;14(4):535–548.

17. Kovács M, Tóth J, Hetényi C, Málnási-Csizmadia A, Sellers JR. Mechanism of blebbistatin inhibition of myosin II. J Biol Chem. 2004;279(34):35557–35563.

18. Hartman EJ, Omura S, Laposata M. Triacsin C: a differential inhibitor of arachidonoyl-CoA synthetase and nonspecific long chain acyl-CoA synthetase. Prostaglandins. 1989;37(6):655–671.

19. Wang Y, Hayes V, Jarocha D, et al. Comparative analysis of human ex vivo-generated platelets vs megakaryocyte-generated platelets in mice: a cautionary tale. Blood. 2015;125(23):3627–3636.

20. Begum J, Day W, Henderson C, et al. A method for evaluating the use of fluorescent dyes to track proliferation in cell lines by dye dilution. Cytometry A. 2013;83A:1085–1095.

21. Barrachina MN, Pernes G, Becker IC, et al. Efficient megakaryopoiesis and platelet production require phospholipid remodeling and PUFA uptake through CD36. Nat Cardiovasc Res. 2023;2(8):746–763.

22. Nakamura-Ishizu A, Takubo K, Fujioka M, Suda T. Megakaryocytes are essential for HSC quiescence through the production of thrombopoietin. Biochem Biophys Res Commun. 2014;454(2):353–357.

23. Fox, J. Applied Regression Analysis and Generalized Linear Models. McMaster University, Canada; 2015.

24. Wickham H. ggplot2: Elegant Graphics for Data Analysis. Springer Link; 2009.

25. Koupenova M, Livada AC, Morrell CN. Platelet and Megakaryocyte Roles in Innate and Adaptive Immunity. Circ Res. 2022;130(2):288–308.

26. Guo L, Shen S, Rowley J, et al. Platelet MHC class I mediates CD8+ T-cell suppression during sepsis. Blood. 2021;138(5):401–416.

27. Zufferey A, Speck ER, Machlus KR, et al. Mature murine megakaryocytes present antigen-MHC class I molecules to T cells and transfer them to platelets. Blood Adv. 2017;1(20):1773–1785.

28. Friedman AH, Fahey JT. The transition from fetal to neonatal circulation: normal responses and implications for infants with heart disease. Semin Perinatol. 1993;17(2):106–121.

29. Davenport P, Liu SZ-J, Sola-Visner M. Fetal vs adult megakaryopoiesis. Blood. 2022;139(22):3233–3244.

30. Takayama N, Eto K. In vitro generation of megakaryocytes and platelets from human embryonic stem cells and induced pluripotent stem cells. Methods Mol Biol. 2012;788:205–217.

31. Nakamura S, Takayama N, Hirata S, et al. Expandable megakaryocyte cell lines enable clinically applicable generation of platelets from human induced pluripotent stem cells. Cell Stem Cell. 2014;14(4):535–548.

32. Sugimoto N, Kanda J, Nakamura S, et al. iPLAT1: the first-in-human clinical trial of iPSC-derived platelets as a phase 1 autologous transfusion study. Blood. 2022;140(22):2398–2402.

33. Asquith NL, Carminita E, Camacho V, et al. The bone marrow is the primary site of thrombopoiesis. Blood. 2024;143(3):272–278.

34. Shin JW, Buxboim A, Spinler KR, et al. Contractile forces sustain and polarize hematopoiesis from stem and progenitor cells. Cell Stem Cell. 2014;14(1):81–93.

35. Barrachina MN, Pernes G, Becker IC, et al. Efficient megakaryopoiesis and platelet production require phospholipid remodeling and PUFA uptake through CD36. Nat Cardiovasc Res. 2023;2(8):746–763.

36. Kyriakides TR, Rojnuckarin P, Reidy MA, et al. Megakaryocytes require thrombospondin-2 for normal platelet formation and function. Blood. 2003;101(10):3915–3923.

37. De Botton S, Sabri S, Daugas E, et al. Platelet formation is the consequence of caspase activation within megakaryocytes. Blood. 2002;100(4):1310–1317.

38. Tiwari S, Italiano JE Jr, Barral DC, et al. A role for Rab27b in NF-E2-dependent pathways of platelet formation. Blood. 2003;102(12):3970–3979.

39. Strassel C, Eckly A, Léon C, et al. Intrinsic impaired proplatelet formation and microtubule coil assembly of megakaryocytes in a mouse model of Bernard-Soulier syndrome. Haematologica. 2009;94(6):800–810.

40. Pecci A, Malara A, Badalucco S, et al. Megakaryocytes of patients with MYH9-related thrombocytopenia present an altered proplatelet formation. Thromb Haemost. 2009;102(1):90–96.

41. Nurden AT, Federici AB, Nurden P. Altered megakaryocytopoiesis in von Willebrand type 2B disease. J Thromb Haemost. 2009;7 Suppl 1:277–281.

42. Evans AL, Dalby A, Foster HR, et al. Transfer to the clinic: refining forward programming of hPSCs to megakaryocytes for platelet production in bioreactors. Blood Adv. 2021;5(7):1977–1990.

43. Feng Q, Shabrani N, Thon JN, et al. Scalable generation of universal platelets from human induced pluripotent stem cells. Stem Cell Reports. 2014;3(5):817–831.

44. Lefrançais E, Looney MR. Platelet Biogenesis in the Lung Circulation. Physiology (Bethesda). 2019;34(6):392–401.

45. Tang L, Liu C, Rosenberger P. Platelet formation and activation are influenced by neuronal guidance proteins. Front Immunol. 2023;14:1206906.

46. Borst S, Sim X, Poncz M, French DL, Gadue P. Induced Pluripotent Stem Cell-Derived Megakaryocytes and Platelets for Disease Modeling and Future Clinical Applications. Arterioscler Thromb Vasc Biol. 2017;37(11):2007–2013.

47. Schmitt A, Guichard J, Massé JM, Debili N, Cramer EM. Of mice and men: comparison of the ultrastructure of megakaryocytes and platelets. Exp Hematol. 2001;29(11):1295–1302.

48. Zauli G, Vitale M, Falcieri E, et al. In vitro senescence and apoptotic cell death of human megakaryocytes. Blood. 1997;90(6):2234–2243.

49. Thon JN, Italiano JE Jr. Does size matter in platelet production? Blood. 2012;120(8):1552–61.

50. Ng WH, Varghese B, Ren X. Understanding and Engineering the Pulmonary Vasculature. Adv Exp Med Biol. 2023;1413:247–264.

51. Jin E, Kawanami O. Unique distribution of von Willebrand factor and thrombomodulin in endothelial cells of human pulmonary microvessels. J Nippon Med Sch. 2000;67(2):64–65.

52. Dunois-Lardé C, Capron C, Fichelson S, Bauer T, Cramer-Bordé E, Baruch D. Exposure of human megakaryocytes to high shear rates accelerates platelet production. Blood. 2009;114(9):1875–1883.

53. Woods MJ, Landon CR, Greaves M, Trowbridge EA. The placenta: a site of platelet production? Platelets. 1992;3(4):211–215.

54. Lill MC, Perloff JK, Child JS. Pathogenesis of thrombocytopenia in cyanotic congenital heart disease. Am J Cardiol. 2006;98(2):254–258.

55. Tilburg J, Becker IC, Italiano JE. Don’t you forget about me(gakaryocytes). Blood. 2022; 139(22):3245–3254.

56. Sugimoto N, Eto K. Generation and manipulation of human iPSC-derived platelets. Cell Mol Life Sci. 2021;78(7):3385–3401.

